# Horizontal gene transfer-initiated reorganization of lipid metabolism drives lifestyle innovation in a eukaryote

**DOI:** 10.1101/2024.08.21.608818

**Authors:** Bhagyashree Dasari Rao, Elisa Gomez Gil, Maria Peter, Gabor Balogh, Vanessa Nunes, James I. MacRae, Qu Chen, Peter Rosenthal, Snezhana Oliferenko

**Author notes:** Corresponding author: Snezhana Oliferenko. These authors contributed equally (Bhagyashree Dasari Rao, Elisa Gomez Gil).

## Abstract

Horizontal gene transfer is a source of metabolic innovation and adaptation to new environments. Yet, how horizontally transferred metabolic functionalities are integrated into host cell biology remains an open question. Here, we use the fission yeast Schizosaccharomyces japonicus to probe how eukaryotic lipid metabolism is rewired in response to the acquisition of a horizontally transferred squalene-hopene cyclase Shc1. We show that Shc1-dependent production of hopanoids, the structural mimics of eukaryotic sterols, allows S. japonicus to thrive in anoxia, where sterol biosynthesis is not possible. We further demonstrate that glycerophospholipid fatty acyl asymmetry, prevalent in S. japonicus, is crucial for accommodating both sterols and hopanoids in membranes, and explain how Shc1 functions alongside the native sterol biosynthetic pathway to support membrane properties. Through engineering experiments in the sister species S. pombe, which naturally lacks Shc1, we show that the acquisition of Shc1 may entail new physiological traits; however, to maximize Shc1 performance, sterol biosynthesis must be dampened. Our work sheds new light on the mechanisms underlying cellular integration of horizontally transferred genes in eukaryotes and provides broader insights into the evolution of membrane organization and function.

## Introduction

The physicochemical properties of biological membranes are fine-tuned to support cellular and organismal physiology. Deregulation of lipid composition may change membrane physicochemical features, affecting the functions of membrane-associated proteins, cell physiology, and survival^1,2^. Eukaryotic membranes include three major lipid classes: glycerophospholipids (GPLs), sphingolipids and sterols.

Sterols are essential components of canonical eukaryotic membranes, maintaining the structure, fluidity, and permeability of the lipid bilayer^3^. Although the sterol biosynthetic pathway is evolutionarily conserved in eukaryotes, it yields different final products in animals (cholesterol), plants (phytosterols), and fungi (ergosterol).

Crucially, *de novo* sterol biosynthesis requires multiple oxygen-dependent steps and thus, eukaryotic membrane chemistry relies heavily on oxygen availability^4^. However, many eukaryotes inhabit or explore hypoxic and anoxic niches^5^, which presumably entails the evolutionary adaptation of their lipid metabolism to maintain membrane integrity in oxygen-deprived environments.

Unlike the popular model fission yeast *Schizosaccharomyces pombe*, which is an obligate aerobe, its relative *Schizosaccharomyces japonicus* thrives in both aerobic and anaerobic environments^6–8^. The *S. japonicus* lineage has diverged from *S. pombe* approximately 200 million years ago^9^, and has lost the ability to respire oxygen, instead becoming a committed fermenting species^10^. *S. japonicus* also has an expanded optimum temperature range, growing at temperatures up to 42°C, unlike *S. pombe* that is restricted to temperatures below 36°C^8^. *S. japonicus* is a dimorphic organism, capable of burrowing deep into the substrate in its hyphal form^11,12^, presumably reaching beyond the limit of oxygen diffusion. Unlike budding yeast that grow well anaerobically only when provided with ergosterol^13^, S*. japonicus* does not require supplementation for anaerobic growth, suggesting that it has evolved a strategy to circumvent the oxygen demands of lipid biosynthesis.

Many prokaryotes use hopanoids as sterol surrogates to regulate membrane properties under harsh conditions^14,15^. Sterols and hopanoids are derived from the same squalene precursor, but hopanoids are cyclized from squalene in a single, oxygen-independent step catalyzed by a squalene-hopene cyclase (SHC)^15^. Interestingly, *S. japonicus* is known to synthesize hopanoids even in normoxia and upregulate hopanoid synthesis upon oxygen depletion^6^. It was suggested that hopanoids were synthesized by an SHC originating from an *Acetobacter*-related species^6^, although the role of this enzyme in *S. japonicus* physiology has not been tested.

Here, we explore the contribution of the SHC to *S. japonicus* lifestyle and explain how it functions alongside the ergosterol biosynthetic pathway to support membrane properties both in the presence and in the absence of oxygen. Using biophysical approaches, we identify the glycerophospholipid fatty acyl asymmetry as a key feature allowing both “native” and “foreign” triterpenoids to function in *S. japonicus* membranes. Finally, by engineering the related fission yeast *S. pombe*, we explore how hopanoids can be integrated into the logic of ergosterol-rich eukaryotic physiology.

## Results

### Both ergosterol and hopanoids support *S. japonicus* lifestyle

*S. japonicus* genome harbors a horizontally transferred gene encoding a squalene-hopene cyclase protein (SJAG_03360, hereafter referred to as Shc1)^6,16^. To synthesize hopanoids, Shc1 is predicted to use squalene, which is also a precursor for the squalene epoxidase Erg1, a rate-limiting enzyme of ergosterol biosynthesis. Unlike Erg1 and several other enzymes in the ergosterol pathway, which require oxygen for their function, SHCs synthesize hopanoids in an oxygen-independent manner^17^ (Fig. 1a). Shc1 tagged at its endogenous locus with superfolder GFP (Shc1-sfGFP) localized to cellular membranes, including the nuclear envelope (NE), cortical endoplasmic reticulum (ER) and/or the plasma membrane, and the vacuolar membranes (Fig. 1b). Unlike the wild-type or Shc1-sfGFP-expressing cells, *S. japonicus* mutant lacking *shc1* was unable to grow in the absence of oxygen (Fig. 1c), demonstrating the requirement for Shc1 in *S. japonicus* anaerobic growth.

**Figure 1.**
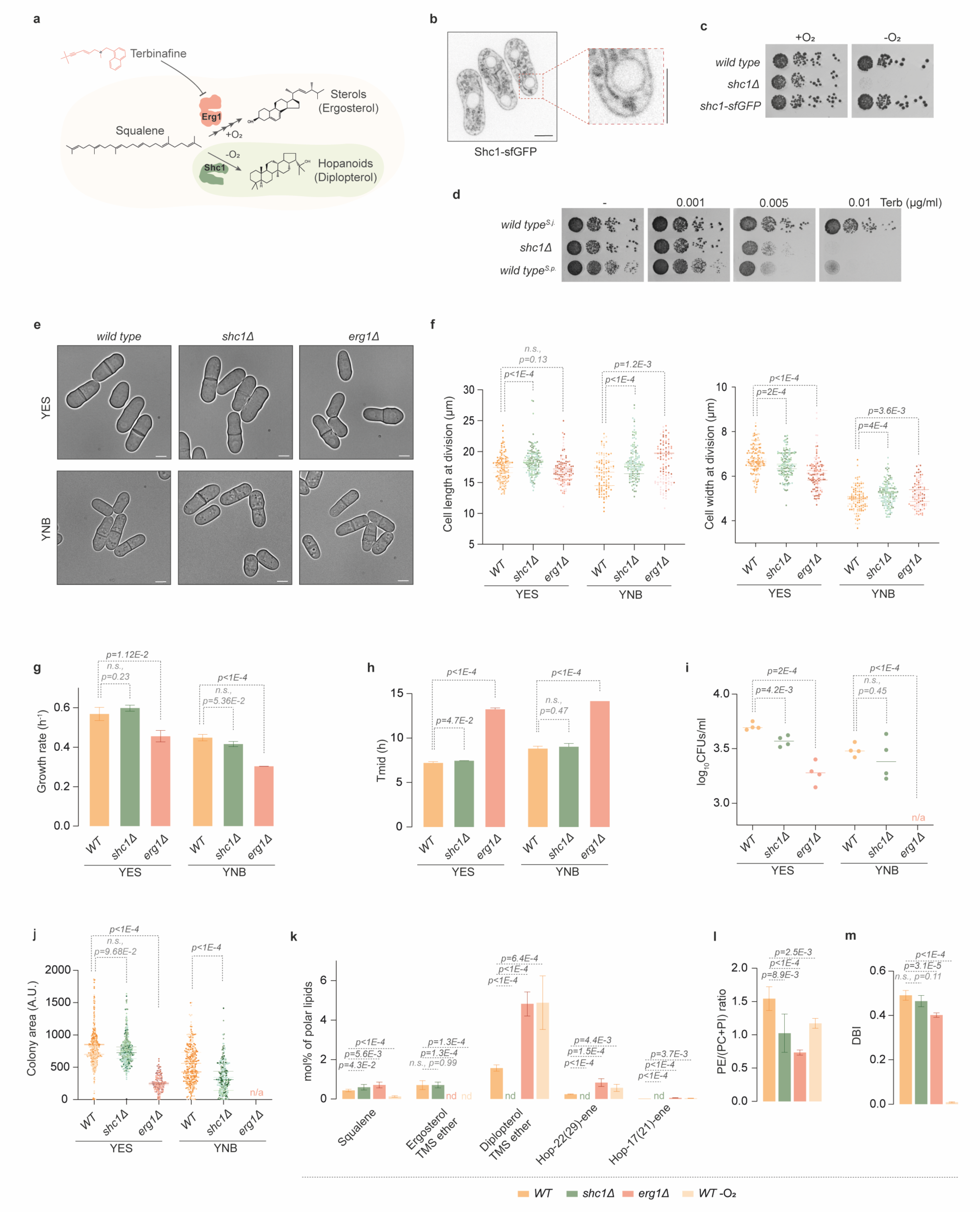
*S. japonicus* relies on hopanoid synthesis to survive in the absence of ergosterol. (**a**) Bifurcation of squalene utilization in *S. japonicus* via 1) multi-step, oxygen-dependent pathway of ergosterol biosynthesis starting with the squalene epoxidase Erg1, or 2) single, oxygen-independent reaction of hopanoid production by the squalene-hopene cyclase Shc1. A selective inhibitor of Erg1, terbinafine, is also shown. (**b**) Single plane spinning disk confocal image of *S. japonicus* cells expressing Shc1-sfGFP grown in YES medium. A magnified image is included. (**c**) Serial dilution assay of *S. japonicus* strains of the indicated genotypes carried out in normoxia or in anoxia in YES medium. (**d**) Serial dilution assay of *S. japonicus wild type* (*WT*) and *shc1Δ* strains, and *S. pombe WT* strain carried out in normoxia in YES medium in the absence or presence of terbinafine (Terb). (**e**) Micrographs of *WT*, *shc1Δ* and *erg1Δ S. japonicus* cells grown in YES or the modified YNB media. (**f**) Quantification of cell length (*left panel*) and width (*right panel*) at division in *S. japonicus WT* (n = 51, 51 and 49 cells grown in YES; n = 44, 36 and 30 cells grown in YNB), *shc1Δ* (n = 54, 50 and 51 cells grown in YES; n = 34, 42 and 66 cells grown in YNB) and *erg1Δ* (n = 52, 34 and 43 cells grown in YES; n = 26, 33 and 45 cells grown in YNB) cultures. Bars represent medians of biological repeats, n = 3. (**g**) Comparison of growth rates of *S. japonicus WT*, *shc1Δ* and *erg1Δ* cells grown in YES or YNB. (**h**) Comparison of the time at which the population reaches ½ of the maximum possible population (Tmid) in the experiments shown in (**g**). (**i**) Survival of YES- or YNB-grown *S. japonicus WT*, *shc1Δ* and *erg1Δ* cells in stationary phase. Colony forming units (CFUs) per ml were counted after 48 h of growth on YES, and 72 h for YNB. (**j**) Colony area quantification (expressed as arbitrary units) of CFUs counted in experiment shown in (**i**). (**k**) GC-MS quantification of triterpenoid content in *S. japonicus WT*, *shc1Δ* and *erg1Δ* cells grown in normoxia, and *WT* cells grown in anoxia. nd, not detected. (**l**) Ratio between phosphatidylethanolamine (PE) and the sum of phosphatidylcholine (PC) and phosphatidylinositol (PI) (PE/(PC+PI)) calculated from the ESI-MS lipidomics data shown in Fig. S1f. (**m**) Comparison of double bond index (DBI) calculated for the four main glycerophospholipid (GPL) classes (PC, PI, PE and phosphatidylserine (PS)) in *S. japonicus WT*, *shc1Δ* and *erg1Δ* cells grown in normoxia, and *WT* grown in anoxia. (**b**, **e**) Scale bars represent 5 µm. (**g**, **h**) Data are represented as average ± S.D. (n=3). (**i**, **j**) Bars represent medians (n=4). (**k**-**m**) Data are represented as average ± S.D. (n=5). (**f**-**m**) p-values are derived from two-tailed unpaired t-test.

Sterols are thought to be crucial for the maintenance of biophysical properties of eukaryotic lipid bilayers^18^. To probe whether hopanoids can act as ergosterol substitutes in *S. japonicus* membranes in normoxia, we treated cells with the Erg1 inhibitor terbinafine, which disrupts sterol biosynthesis^19^ (Fig. 1a). Contrary to the *S. japonicus* wild type, the *shc1Δ* mutant exhibited a marked growth sensitivity to terbinafine. As expected, a related species *S. pombe*, which does not harbor an SHC^9^, was as sensitive to the drug as the *S. japonicus shc1Δ* strain (Fig. 1d). We concluded that *S. japonicus* relies on Shc1 activity when ergosterol biosynthesis is inhibited.

Since sterols are critical components of canonical eukaryotic membranes, the *erg1* gene is essential for life in *S. pombe* and other fungi^20–22^. Strikingly, we were able to generate *S. japonicus erg1Δ* mutant, suggesting that this fission yeast does not strictly require sterols to support its growth (Fig. 1e). As expected, *erg1Δ* cells were not stained by the naturally fluorescent sterol marker polyene filipin^23^ (Fig. S1a), and exhibited resistance to amphotericin B, a drug that induces the formation of pores in the plasma membrane by binding ergosterol^24^ (Fig. S1b).

To assess how Shc1 and Erg1 contribute to *S. japonicus* physiology in normoxia, we investigated cellular growth and survival of *shc1Δ* and *erg1Δ* mutants both in the rich Yeast Extract with Supplements (YES) and in the modified minimal Yeast Nitrogen Base (YNB) media (see Materials and Methods). Fission yeast cells grow at cell tips and divide in the middle, exhibiting stereotypic pill-shaped geometry in exponentially growing cultures^25^. In minimal medium, where the anabolic demands are high, *S. japonicus* grows slower and undergoes pronounced downscaling of its geometry, dividing at decreased length and width but maintaining its aspect ratio^26^. *S. japonicus* cells lacking *shc1* exhibited somewhat perturbed geometry, dividing at increased cell length. Cell width was also deregulated (Fig. 1e, f). Despite differences in cell geometry, *shc1Δ* cultures grew at normal rates at 30°C (Fig. 1g, h). Interestingly, *shc1Δ* cells showed decreased viability at 40°C in the minimal medium (Fig. S1c). Additionally, cells lacking Shc1 exhibited mild defects in survival and regrowth from the stationary phase (Fig. 1i, j and Fig. S1d).

The lack of Erg1 led to more pronounced phenotypes. At the optimal temperature of 30°C, *erg1Δ* cells exhibited defects in cell geometry (Fig. 1e, f), slower growth rate and extended lag-phase (Fig. 1g, h). Cells lacking Erg1 were less viable at higher temperatures in the minimal medium (Fig. S1c). Cell survival in stationary phase and subsequent regrowth were strongly affected by the absence of sterols, with the *erg1Δ S. japonicus* mutant not being able to form colonies in the minimal medium (Fig. 1i, j and Fig. S1d). Thus, it appears the aerobic physiology of *S. japonicus* relies in larger part on Erg1 rather than Shc1.

To understand how the lack of Shc1 or Erg1 impacted membrane lipid composition, we performed gas chromatography-mass spectrometry (GC-MS) and shotgun electrospray ionization mass spectrometry (ESI-MS) analyses of *S. japonicus* total lipid extracts. As previously described^6^, *S. japonicus* wild-type cells produced high levels of hopanoids even in the presence of oxygen, with diplopterol being the most abundant. We also detected the hopenes hop-22(29)-ene and hop-17(21)-ene (Fig. 1k and Data S1, Table 1).

The *S. japonicus shc1Δ* mutant did not produce hopanoids, demonstrating that Shc1 is indeed responsible for hopanoid synthesis in this organism. As expected, we did not detect ergosterol in *erg1Δ* cells grown in normoxia or in the wild-type cells grown in anaerobic environment. Interestingly, whereas the cellular amount of hopanoids increased profoundly when ergosterol synthesis was inhibited either by *erg1Δ* mutation or the growth in anoxia, the converse was not true. Cells lacking Shc1 and wild-type cells exhibited comparable ergosterol content (Fig. 1k and Data S1, Table 1). The amount of squalene, the substrate for both Shc1 and Erg1, dropped dramatically in the absence of oxygen (Fig. 1k).

Notably, the lack of either hopanoids or ergosterol led to changes in the total cellular lipid landscape, as shown by the SoamD score^27^ (Fig. S1e). In cells lacking Shc1, the ratio between phosphatidylethanolamine (PE) and the sum of phosphatidylcholine and phosphatidylinositol (PE/(PC+PI)) was significantly reduced, suggesting a cellular response to stabilize the membranes^28^ (Fig. 1l and Fig. S1f). We also detected decreased abundance of lysophosphatidylethanolamine (LPE) and ceramide (Cer), and increased levels of phosphatidic acid (PA), diacylglycerol (DG) and inositol phosphoceramide (IPC) (Fig. S1g).

The cellular PE/(PC+PI) ratio was similarly decreased when cells were unable to synthetize sterols, either in the *erg1Δ* mutant or when grown in the absence of oxygen (Fig. 1l and Fig. S1f). In addition, the level of FA chain desaturation, already low in *S. japonicus*^29^, decreased further in *erg1Δ* mutant (Fig. 1m and Fig. S1i, j), and the average FA chain length mildly increased (Fig. S1h). The decrease in PE/(PC+PI) ratio, decrease in FA desaturation and an increase in GPL chain length might all contribute to membrane stability in the absence of sterols^28^. As expected, FA desaturation was virtually absent in anaerobic conditions, due to the oxygen dependence of the delta-9 desaturase Ole1^30^, with cells producing higher amounts of asymmetrical GPL species (Fig. 1m and Fig. S1i, j). We also detected differences in the abundance of some of the minor GPL and lysoglycerophospholipid classes, sphingolipids, and storage lipids in both *erg1Δ* mutant and the wild type growing in anoxia (Fig. S1g and Data S1, Table 1).

Taken together, our results suggest that ergosterol and hopanoids collaborate to support membrane properties in *S. japonicus* in a variety of physiological situations. Importantly, the overall lipidome of this organism has been adapted to the acquisition of hopanoid biosynthesis through horizontal gene transfer.

### Asymmetrical saturated lipids can use either hopanoids or ergosterol to support membrane properties

To understand how *S. japonicus* may rely on either ergosterol or hopanoids to support its physiology, we turned to a bottom-up approach relying on model membranes. We used PC(C18:0/C10:0) (1-stearoyl-2-decanoyl-sn-glycero-3-phosphocholine, SDPC) and either PC(C16:0/C18:1) (1-palmitoyl-2-oleoyl-sn-glycero-3-phosphocholine, POPC) or PC(C18:1/C18:1) (1,2-dioleoyl-sn-glycero-3-phosphocholine, DOPC) as models for asymmetrical saturated *S. japonicus*- or symmetrical unsaturated *S. pombe*-like glycerophospholipids, respectively (Fig. 2a and ^29^).

**Figure 2.**
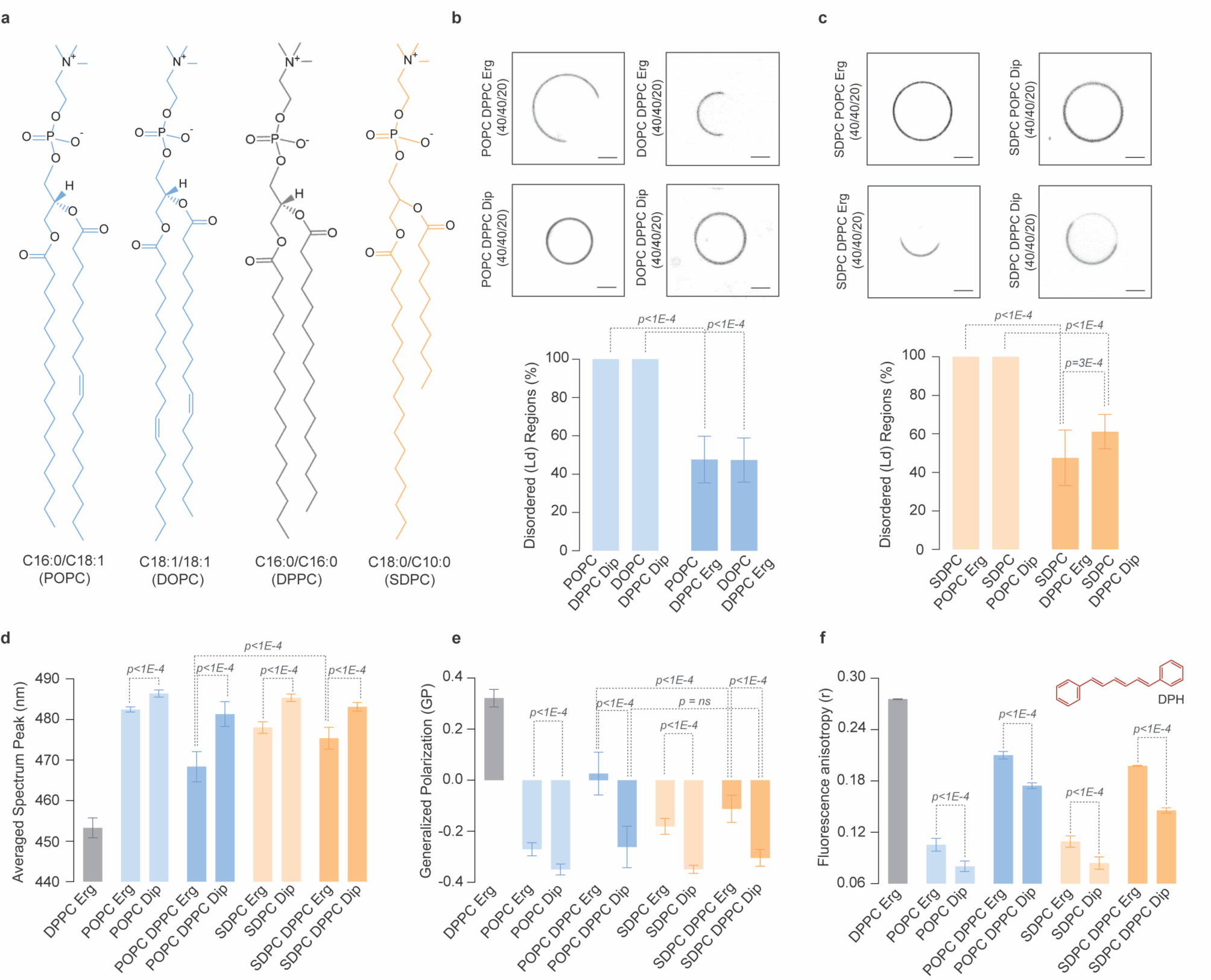
Asymmetrical glycerophospholipids support membrane phase properties both in the presence of ergosterol and diplopterol. (**a**) Chemical structures of glycerophospholipids used in the study. (**b**, **c**) The top panel shows representative spinning disk confocal images (*mid-plane*) of three-component GUVs assembled with equimolar amounts of symmetrical unsaturated glycerophospholipids POPC or DOPC and the gel-like DPPC (**b**), or asymmetrical saturated glycerophospholipid SDPC (**c**) with either 20 mol% ergosterol or diplopterol. GUVs were labelled with FAST DiI, a marker for disordered membrane regions. Bar charts depict the quantification of area of liquid-disordered (Ld) regions upon phase separation (average ± S.D. from individual measurements of at least 50 GUVs). (**d**) Estimation of membrane polarity from spectral GP imaging. Average spectrum peaks shown for two- and three-component GUVs made with the symmetrical unsaturated lipid POPC or the asymmetrical saturated lipid SDPC. Peak values were estimated from the normalized intensity vs wavelength plots (Fig. S2g, h) of individual measurements (average ± S.D. obtained from at least 15 GUVs). (**e**) Quantification of membrane order in two- and three-component GUVs made with the symmetrical unsaturated lipid POPC or the asymmetrical saturated lipid SDPC, inferred from (**d**). GP values were calculated using Equation (i) (n = at least 15, see Materials and Methods). (**f**) Membrane order measured by DPH fluorescence anisotropy (*structure shown in top right corner*) in two- and three component LUVs made with the symmetrical unsaturated POPC or the asymmetrical saturated SDPC (average ± S.D. of at least three independent measurements). (**b**-**f**) p-values obtained by unpaired parametric t-test. (**b**, **c**) Scale bars represent 2 μm.

When combined with 30 mol% of either ergosterol or diplopterol, all three glycerophospholipids formed disordered membranes, as indicated by the labelling of giant unilamellar vesicles (GUVs) with FASTDiI^31,32^, a dye that partitions preferentially into the liquid-disordered (Ld) membrane phase (Fig. S2a, b). Suggesting lower membrane order, membranes formed with the asymmetrical glycerophospholipid SDPC in single or two-component mixtures with either triterpenoid exhibited higher water permeability than those made with the symmetrical unsaturated POPC (Fig. S2c, d).

When mixed with the gel-forming symmetrical saturated glycerophospholipid PC(C16:0/C16:0) (1,2-dipalmitoyl-sn-glycero-3-phosphocholine, DPPC) and sterols, the unsaturated GPLs tend to separate into the liquid-disordered (Ld) and liquid-ordered (Lo) phases, recapitulating a key property of biological membranes^47^. Importantly, both *S. pombe*-like symmetrical unsaturated glycerophospholipids, POPC or DOPC, exhibited phase separation in the presence of 20 mol% ergosterol but not diplopterol (Fig. 2b).

We then explored the phase behavior in three-component liposomes containing the *S. japonicus*-like asymmetrical SDPC glycerophospholipid. Confirming that it conferred disorder to membranes (Fig. S2b-d), SDPC did not support phase separation when combined with the unsaturated symmetrical POPC. Strikingly, either 20 mol% ergosterol or 20 mol% diplopterol was sufficient to induce phase separation in liposomes containing SDPC together with the gel-forming DPPC (Fig. 2c). Of note, ergosterol was more efficient in promoting phase separation in membranes containing the asymmetrical GPL (Fig. 2c).

Our results so far indicated that asymmetrical saturated lipids are highly disordered but can accommodate either ergosterol or diplopterol to promote phase separation in membranes. To further explore the properties of such membranes, and to compare them with membranes made from more conventional, symmetrical unsaturated GPLs, we estimated membrane order in GUVs using the fluorescent probe C-laurdan (structure in inset of Fig. S2e). The emission properties of this probe depend on the lipid bilayer environment and can be used to estimate relative levels of lipid packing by calculating a parameter called Generalized Polarization (GP)^33–35^.

The membranes composed of the gel-forming DPPC and ergosterol displayed the maximum blue-shifted peak (∼453 nm) and the highest GP, indicating that they were indeed highly ordered (Fig. 2d, e, and Fig. S2e, f). In two-component mixtures containing either symmetrical unsaturated POPC or asymmetrical saturated SDPC, ergosterol supported higher membrane order as compared to diplopterol. In the case of *S. pombe*-like three-component membranes, the phase-separated GUVs made of POPC, DPPC and ergosterol showed significantly higher membrane order as compared to the non-phase-separating GUVs made with diplopterol. Interestingly, even though both ergosterol- and diplopterol-containing three-component SDPC-based membranes showed phase separation (Fig. 2c), the overall membrane order was lower in the presence of diplopterol (Fig. 2d, e, and Fig. S2e, f).

To corroborate our C-laurdan imaging data, we measured the anisotropy of the rod-shaped fluorophore 1,6-diphenyl-1,3,5-hexatriene (DPH) (structure in inset of Fig. 2f). DPH fluorescence anisotropy depends on its rotational mobility in the bilayer. It is more restricted in ordered environments, producing higher anisotropy values^36,37^. As expected, the control gel-like large unilamellar vesicles (LUVs) made of DPPC and ergosterol showed the highest anisotropy. Similarly to C-laurdan measurements, ergosterol promoted higher order in the context of all two- and three-component lipid mixtures, as compared to diplopterol (Fig. 2f).

Overall, our model membrane data suggest that ergosterol is more efficient than diplopterol at promoting membrane order. However, both triterpenoids can facilitate phase separation in membranes containing the asymmetrical saturated glycerophospholipids abundant in *S. japonicus*. In contrast, only ergosterol can achieve this crucial effect in membranes composed of symmetrical *S. pombe*-like glycerophospholipids.

### Cryo-EM imaging demonstrates that asymmetrical glycerophospholipids support membrane phase separation in the presence of either ergosterol or diplopterol

To directly image our model membranes without introducing external probes, we employed cryogenic electron microscopy (cryo-EM), which has been successfully used to detect nanoscopic domains in synthetic and bioderived membranes^38,39^. Membrane thickness can be estimated by D_TT_, a distance between two troughs in intensity across the bilayer, which correspond to electron-rich head group regions in glycerophospholipids. Strikingly, single-component membranes made of saturated asymmetrical SDPC were considerably thinner (D_TT_ = 27.78 nm ± 1.72 nm, n = 20) as compared to those composed of symmetrical unsaturated POPC (D_TT_ = 36.39 nm ± 2.83 nm, n = 20) (Fig. 3a, b).

**Figure 3.**
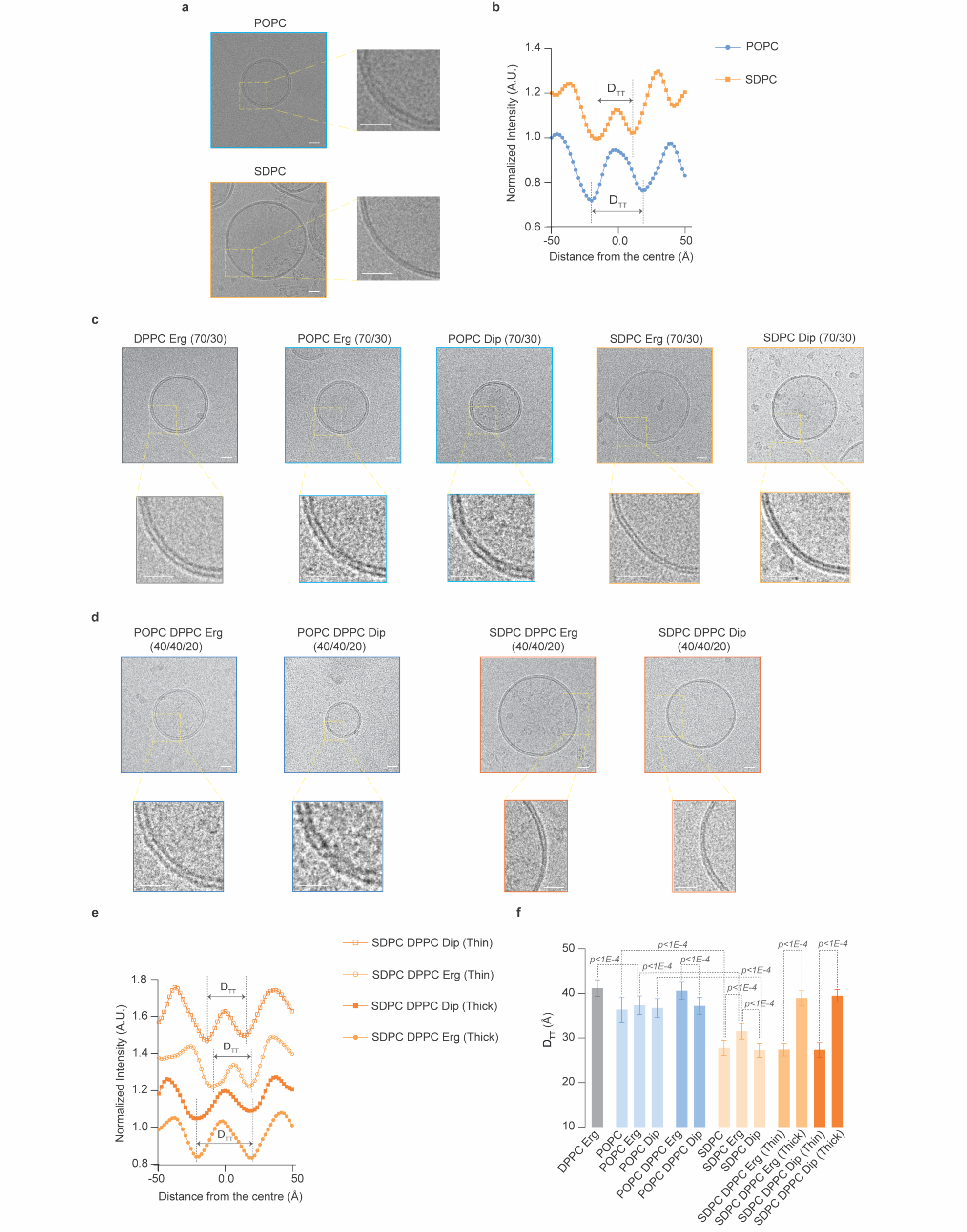
Cryo-EM measurements show that asymmetrical glycerophospholipids form thinner membranes and support phase separation with either ergosterol or diplopterol. (**a**) Representative cryo-EM images of single component LUVs made from the monounsaturated symmetrical lipid POPC (*top panel*) and the asymmetrical saturated SDPC (*bottom panel*). *Right*, magnified areas of respective images. (**b**) Representative normalized intensity profiles for POPC and SDPC liposomes. The trough-to-trough distance D_TT_ is shown by arrows within dotted lines. (**c**) Representative cryo-EM images of two-component liposomes made from either POPC or SDPC glycerophospholipids in combination with 30 mol% ergosterol or diplopterol. The gel-like membranes composed of the symmetrical saturated lipid DPPC with ergosterol is also included. Magnified areas are shown at the bottom of respective images. (**d**) Representative cryo-EM images of three-component liposomes made either from 40 mol% POPC or SDPC glycerophospholipids in combination with 40 mol% gel-forming DPPC and 20 mol% ergosterol or diplopterol. Magnified areas are shown at the bottom of respective images. Note the phases of different membrane thickness in the ternary SDPC-DPPC-based membranes. (**e**) Representative normalized intensity profiles for “thick” and “thin” regions in phase-separated liposomes made from the asymmetrical saturated lipid SDPC in the presence of DPPC and ergosterol or diplopterol. The trough-to-trough distance D_TT_ is indicated by arrows. (**f**) The graph shows D_TT_ values for single, two- and three-component LUVs assembled with POPC or SDPC (average ± S.D. n= 20 liposomes, 10 measurements for each liposome). The p-values were obtained from unpaired parametric t-tests. (**a**, **c**, **d**) Scale bars represent 20 nm.

We observed a similar trend in two-component lipid mixtures. Whereas the control gel-like liposomes (DPPC with ergosterol) exhibited the highest D_TT_, both ergosterol- and diplopterol-containing SDPC bilayers were thinner than their counterparts made of symmetrical unsaturated POPC. Consistent with its greater ordering potential, ergosterol increased the thickness of SDPC-containing membranes (Fig. 3c, see quantification in Fig. 3f and Fig. S3).

Using cryo-EM, we were not able to detect coexisting membrane domains in three-component phase-separated *S. pombe*-like liposomes (POPC/DPPC/ergosterol), likely due to relatively small differences in membrane thickness between DPPC and POPC (Fig. 3d, quantification in Fig. 3f and Fig. S3). Remarkably, we observed a clear phase separation within single liposomes in three-component mixtures made with asymmetrical *S. japonicus*-like SDPC glycerophospholipid (Fig. 3d-f). The width measurements of “thick” (ordered) and “thin” (disordered) domains were comparable in SDPC membranes containing either ergosterol or diplopterol (Fig. 3d-f).

To conclude, our EM data indicate that (1) asymmetrical saturated glycerophospholipids form thinner membranes; and (2) they can indeed phase separate in the presence of a gel-forming lipid and either ergosterol or diplopterol. The remarkably similar hydrophobic thickness values for the ordered and disordered domains in asymmetrical saturated glycerophospholipid-based membranes containing either ergosterol or diplopterol suggest that these membranes could accommodate similar sets of proteins and *in vivo* functions, regardless of the triterpenoid type.

### Hopanoid synthesis in *S. pombe* enables new physiological features

Our results so far suggested that eukaryotic lipidomes must be adapted to accommodate the bacterial sterol mimics hopanoids. *S. japonicus* appears to solve this problem at least in part by maintaining high levels of FA asymmetry in glycerophospholipids (Fig. 2 and 3), which is a minor feature in the lipidomes of its relative *S. pombe* and budding yeast^29,40^. We wondered if introducing Shc1 into a “naïve” organism, such as the sister species *S. pombe*, could lead to changes in cellular lipid landscape and provide immediate benefits for adapting to new environments.

To this end, we integrated a construct encoding the *S. japonicus* Shc1 tagged with sfGFP as a single copy under the control of a strong, constitutive *tdh1* promoter (*ptdh1:shc1^S.j^.-sfGFP*) in the *S. pombe* genome. In *S. pombe*, Shc1^S.j.^-sfGFP localized predominantly to the NE and cortical ER and/or plasma membrane, reminiscent of its localization in *S. japonicus* (Fig. 4a). Triterpenoid analysis by GC-MS showed that *S. pombe* cells expressing Shc1^S.j.^ synthesized diplopterol (Fig. 4b), although its total cellular amount was considerably lower than in *S. japonicus* (0.134 ± 0.017 mol% polar lipids in *S. pombe* vs 1.567 ± 0.156 in *S. japonicus*). We did not detect two minor hopanoid species present in *S. japonicus* – hop-22(29)-ene and hop-17(21)-ene – in Shc1^S.j.^-expressing *S. pombe* (Data S1, Table 2).

**Figure 4.**
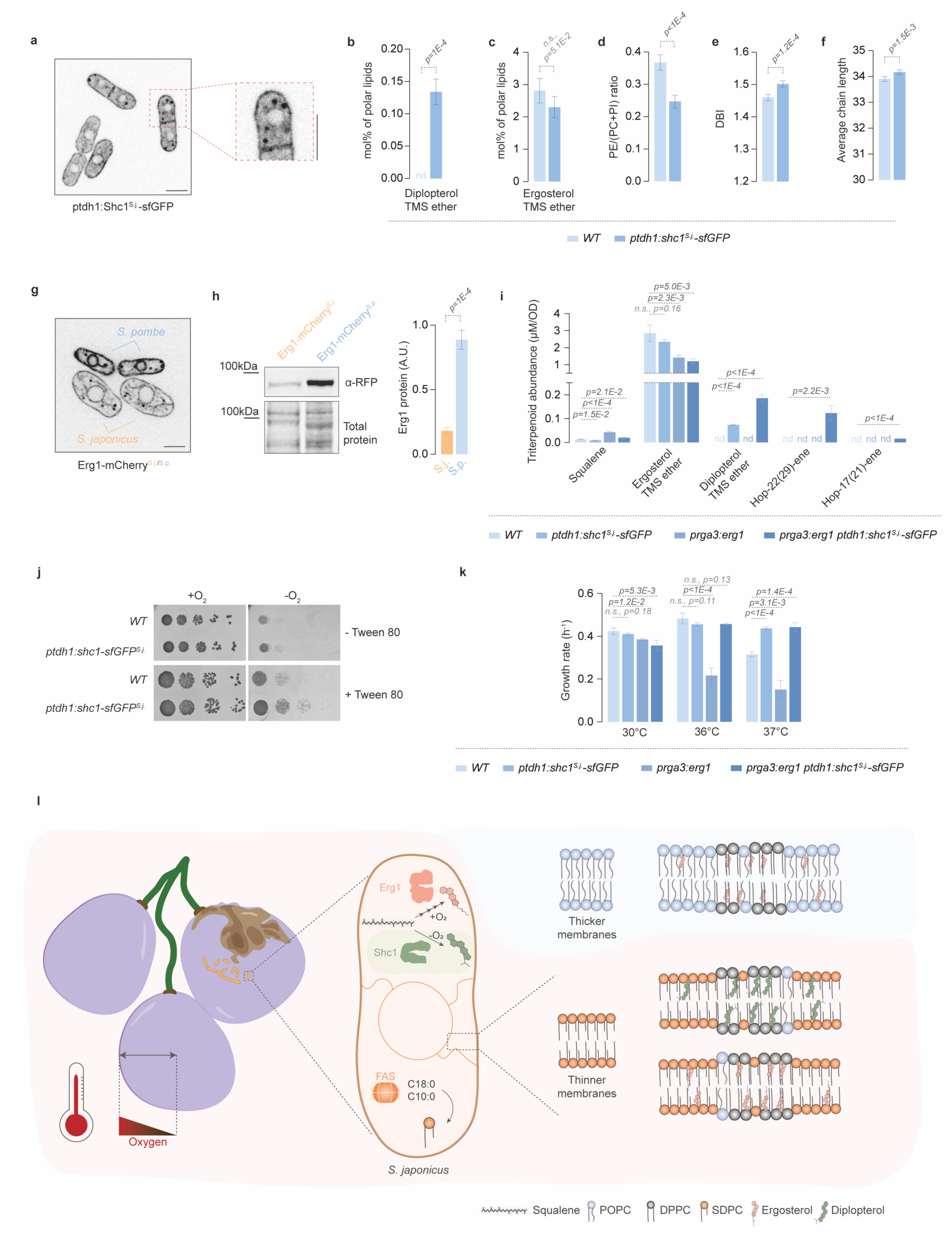
Engineering of hopanoid production in *S. pombe* offers physiological advantages but their efficient synthesis requires dampening of Erg1 levels. (**a**) Single plane spinning disk confocal image of *S. pombe* cells expressing Shc1-sfGFP under the regulation of *tdh1* promoter (*ptdh1:shc1^S.j.^-sfGFP*) grown in YES medium. A magnified image is included. (**b**, **c**) Relative abundance of diplopterol (**b**) and ergosterol (**c**) in *S. pombe wild type* (*WT*) and *ptdh1:shc1^S.j.^-sfGFP* cells. (**d**) PE/(PC+PI) ratio calculated from data shown in Fig. S4a for *S. pombe WT* and *ptdh1:shc1^S.j.^-sfGFP* cells. (**e**) Comparison of the double bond indexes (DBI) calculated for the four main GPL classes in *S. pombe WT* and *ptdh1:shc1^S.j.^-sfGFP* cells. (**f**) Average combined FA length calculated for the sum of PC, PI, PE, and PS in the two strains. (**g**) Single plane spinning disk confocal image of *S. japonicus* and *S. pombe* cells expressing Erg1-mCherry grown in YES. (**h**) Western blot of Erg1-mCherry in the strains shown in (**g**). Revert 700 stain was used to visualize total protein loading. The right panel shows the quantification, expressed as arbitrary units. Data are represented as average ± S.D. (n=3). (**i**) Abundance of triterpenoids in *S. pombe* strains of indicated genotypes, as detected by GC-MS (in µM/OD). Results are represented as average ± S.D. (n=4). (**j**) Serial dilution assay of *S. pombe* strains of indicated genotypes carried out in normoxia or anoxia in YES in the absence or the presence of an unsaturated FA supplement Tween 80. (**k**) Comparison of growth rates of *S. pombe* strains of indicated genotypes. Cells were grown in YES at the indicated temperatures. Data are represented as average ± S.D. (n=3). (**l**) A model suggesting how the acquisition of a squalene-hopene cyclase Shc1 through horizontal transfer has led to the reorganization of *S. japonicus* lipid metabolism, potentially allowing it to explore new ecological niches. The pictorial legend for lipids is included. (**a**, **g**) Scale bars represent 5 µm. (**b**, **i**) nd, not detected. (**b**-**f**) Data are represented as average ± S.D. (n=5). (**b**-**f**, **h**, **i**, **k**) p-values are derived from two-tailed unpaired t-test.

Enabling hopanoid synthesis in *S. pombe* significantly altered the overall cellular lipidome. Although ergosterol levels were not greatly affected (Fig. 4c), we observed differences in the abundance of major GPL classes (Fig. S4a), resulting in a significant decrease in the cellular PE/(PC+PI) ratio (Fig. 4d). We also detected major changes in the abundance of lysoglycerophospholipids, sphingolipids and storage lipids in *ptdh1:shc1^S.j.^-sfGFP S. pombe* cells as compared to the wild type (Fig. S4b).

Membrane glycerophospholipids in *S. pombe* largely consist of symmetrical di- and mono-unsaturated species [36:2] and [34:1], although *S. pombe* also synthesizes some asymmetrical saturated [26:0] and [28:0] GPLs^29,40^. Interestingly, we observed an increase in the [36:2] GPLs with a concomitant decrease in [26:0] and [28:0] species in Shc1^S.j.^-expressing cells (Fig. S4c). Overall, *S. pombe* cells expressing Shc1^S.j.^ showed higher FA desaturation and an increase in average FA chain length (Fig. 4e, f, and Fig. S4d, Data S1, Table 2).

We wondered why the *S. japonicus* Shc1, even when expressed at high levels in *S. pombe*, synthesized only limited amounts of its product diplopterol. Considering that both Shc1 and the squalene epoxidase Erg1 use the same squalene substrate (Fig. 1a), we wondered if competition from native Erg1 could interfere with efficient squalene utilization by Shc1. Consistent with this possibility, Erg1 was much more abundant in *S. pombe* than in *S. japonicus* (Fig. 4g, h). We reasoned that decreasing Erg1 abundance in *S. pombe* might promote the flux of squalene towards hopanoid synthesis. To test this hypothesis, we generated *S. pombe* strains in which the endogenous *erg1* promoter was replaced by the mild constitutive *rga3* promoter^41^ (*prga3:erg1*), either on its own or in combination with *ptdh1:shc1^S.j.^-sfGFP*. Reverse-transcription qPCR analysis showed that the steady-state abundance of *erg1* mRNA in the *prga3:erg1* cells decreased to ∼50% of the wild-type level (Fig. S4e).

The GC-MS analysis confirmed that the ergosterol content was indeed lower in the strains with attenuated expression of *erg1* (Fig. 4i, Data S1, Table 3). Interestingly, we observed a considerable increase in squalene levels in the *prga3:erg1* cells. Shc1^S.j.^ expression in this mutant background led to the reduction in squalene, suggesting that the two enzymes indeed compete for the same substrate pool (Fig. 4i). Further supporting this hypothesis, diplopterol content was ∼2.5 times higher in Shc1^S.j.^-expressing *prga3:erg1* cells, as compared to the wild type with Shc1^S.j.^. Moreover, we also detected the minor products of Shc1^S.j.^, the hop-22(29)-ene and hop-17(21)-ene hopenes, in Shc1^S.j.^-expressing *prga3:erg1* mutant (Fig. 4i). We concluded that the optimization of SHC performance in the eukaryotic context may necessitate tuning the expression and/or activity of the squalene epoxidase.

To illuminate the immediate consequences of introducing hopanoid synthesis in a eukaryote, we tested if *S. pombe* cells expressing Shc1^S.j.^ differed from the wild type in several physiological scenarios, which affect membrane function. Unlike *S. japonicus*, *S. pombe* is an obligate aerobe, which can divide only a few times in fully anoxic conditions, likely due to increasing dilution of sterols, unsaturated FAs and other metabolites that require oxygen for their synthesis (Fig. 4j and ^8^). Strikingly, as long as we supplemented the unsaturated fatty acid Tween 80, the Shc1^S.j.^-expressing *S. pombe* could grow considerably better in the absence of oxygen (Fig. 4j).

*S. pombe* growth is limited at temperatures exceeding 36°C (Fig. 4k and Fig. S4f). Interestingly, although the expression of Shc1^S.j.^ did not affect growth within the physiological range of this organism, it became advantageous at the higher temperature of 37°C (Fig. 4k and Fig. S4f). The beneficial effect of hopanoids on growth at higher temperatures was even more pronounced in cells with attenuated *erg1* expression, which by itself conferred a growth defect at higher temperatures (Fig. 4k and Fig. S4f).

We conclude that acquiring hopanoid biosynthesis through horizontal gene transfer may offer organismal advantages for exploring anoxic and warm ecological niches. Notably, integrating this new module into the metabolism of the recipient cells may require adjustments to the sterol production pathway, alongside molecular adaptations that enhance membrane functionality in the presence of both triterpenoids (Fig. 4l).

## Discussion

Unlike most eukaryotes, the fission yeast *S. japonicus* thrives in strictly anaerobic conditions. Our work suggests that the horizontal acquisition of a bacterial gene encoding a squalene-hopene cyclase^6,9^ has been at the root of this physiological innovation (Fig. 1).

Shc1 appears to be deeply integrated into the lipid metabolism of *S. japonicus*, which produces both hopanoids and ergosterol under normoxic conditions and switches entirely to hopanoids for anaerobic growth. Furthermore, although sterols are typically essential to support membrane function in eukaryotic cells^42^, *S. japonicus erg1Δ* mutant cells, lacking ergosterol production altogether, are viable (Fig. 1e).

*S. japonicus* lipidome is unusually rich in asymmetrical glycerophospholipids containing long (typically C18:0) and medium fatty acyl (C10:0) chains^29^. We show that these lipids form thinner membranes, as compared to those made from the symmetrical unsaturated *S. pombe*-like glycerophospholipid POPC (Fig. 3). This observation potentially explains the shortening of transmembrane helices in a subset of proteins in *S. japonicus* as compared to *S. pombe*^29^, as a strategy to reduce hydrophobic mismatch^43^.

Importantly, we show that when mixed with the gel-like saturated DPPC, these asymmetrical glycerophospholipids support membrane phase separation in the presence of both the native triterpenoid ergosterol and the “foreign” triterpenoid diplopterol (Fig. 2c). This behavior contrasts sharply with symmetrical unsaturated glycerophospholipids DOPC and POPC, which support phase separation only in the presence of ergosterol. It is possible that accommodating bulky diplopterol molecules in the membrane requires the more relaxed lipid packing afforded by relatively compact asymmetrical glycerophospholipids^44^.

Tuning the lipid composition and packing in model membranes may alter phase separation tendencies and properties of phase-separated domains. For instance, diplopterol has been shown to support phase separation in the mixtures with a synthetic sphingolipid and DOPC^47^. Notably, although diplopterol supports membrane ordering in model membranes containing asymmetrical glycerophospholipids, it is a less effective order inducer than ergosterol (Fig. 2d-f). The situation probably differs *in vivo*, given that *S. japonicus* thrives in the absence of oxygen. Interestingly, the lack of Erg1 appears to have some deleterious effects in normoxia, suggesting that ergosterol might have specific functions in aerobic membranes. Alternatively, sterol biosynthesis could function as an oxygen sink, protecting cells from oxidative damage.

Asymmetrical C10:0- and C12:0-containing glycerophospholipids are found in other species, albeit at lower abundances^29,45^. The proportion of such lipids increases in anoxia not only in *S. japonicus* but also budding yeast (Fig. S1i and ^46,47^). This suggests that while the production of the medium-chain FAs is a regulatable feature in many organisms, it might have been constitutively augmented during the evolution of *S. japonicus* following the acquisition of SHC. Indeed, engineering approaches indicate that just a few mutations can change the FAS product spectrum^48^. The ability to produce asymmetrical glycerophospholipids is advantageous not only for accommodating diplopterol but also to maintain membrane fluidity in anoxia where FA desaturation is not possible (Fig. 2 and ^46^).

If oxygen availability is not a consideration, other GPL architectures, such as high proportion of FA desaturation, may possibly support integration of hopanoids^49^. Consistent with this scenario, *S. pombe* cells expressing Shc1^S.j^ further increase FA chain unsaturation rather than synthesizing more asymmetrical glycerophospholipids (Fig. 4e).

Our data suggest that, beyond reducing oxygen dependence, the acquisition of hopanoid biosynthesis may further expand the environmental versatility of unicellular eukaryotes, including their ability to grow at elevated temperatures (Fig. S1c, Fig. 4k, Fig. S4f, and ^8^). Although the molecular mechanisms underlying this effect remain to be elucidated, it is noteworthy that in bacteria, hopanoids have been linked to tolerance to higher temperatures, and acidic and ethanol stresses^50–52^.

Integrating SHC into the logic of ergosterol-reliant membrane lipid metabolism may require cells to co-regulate hopanoid and sterol production. Our data suggest that, at least in fission yeast, Erg1 directly competes with Shc1 for squalene, and thus, its activity must be dampened in order to allow efficient synthesis of hopanoids. Additionally, cells may need to adapt their membrane homeostasis to the presence of different triterpenoids. Accordingly, we observe large-scale changes to the cellular lipidomes both upon the loss (Fig. 1l, Fig. S1e, f, g) and ectopic acquisition of hopanoids (Fig. 4d-f and Fig. S4a-d) in *S. japonicus* and *S. pombe*, respectively.

Sterols are essential in most eukaryotes, and if they cannot be synthesized *de novo*, they must be assimilated from the environment. In such cases, the source of dietary sterols becomes critical. Potential trade-offs of sterol import could be a disruption of native membrane organization and function by “foreign” sterols^53^. Such considerations could drive organisms frequenting hostile environments towards different solutions, including integration of SHC enzymes. In line with this, fission yeasts cannot import sterols^54^. Similar considerations could have been at play in other lineages exhibiting SHC horizontal gene transfer. Of interest, SHCs have been acquired independently in a number of eukaryotic lineages, including several species of pathogenic fungi such as *Aspergillus fumigatus*^55^, which encounter warm hypoxic environments at infection sites^56^.

Horizontal gene transfer is widespread in all major eukaryotic groups and has been linked to metabolic innovations and adaption to new environments^57^. SHC is a standalone metabolic enzyme, which does not require high physical connectivity with other proteins (Fig. 4 and ^58^). This feature, combined with the immediate adaptive benefits of hopanoids in certain environmental conditions (Fig. 4j-l), may explain the multiple independent instances of horizontal gene transfer of SHC and SHC-related enzymes into eukaryotic lineages^59^. Importantly, our work suggests that following the initial acquisition event, the “domestication” of SHC and likely other new metabolic functionalities may necessitate large-scale rewiring of host metabolism with profound consequences for membrane structure and other key aspects of cellular biology.

## Materials and Methods

### Fission yeast strains and growth conditions

The *S. japonicus* and *S. pombe* strains used in this work are listed in Data S2. All strains were prototrophic. Standard fission yeast media and culture methods were used^60–63^, with an exception of the modified minimal yeast nitrogen base (YNB) medium (YNB containing 111 mM glucose, 14.7mM potassium hydrogen phthalate, 15.5mM disodium hydrogen phosphate, and the following supplements: adenine (93.75 mg/l), uracil (75 mg/l), histidine (75 mg/l) and leucine (75 mg/l). We chose a YNB-based minimal medium as an alternative to the canonically-used Edinburgh Minimal Medium (EMM)^60^ since in our experimental setup, *S. japonicus* did not grow in EMM under strictly anaerobic conditions. For liquid cultures, cells were routinely grown in rich yeast extract with supplements (YES) or minimal modified YNB medium in 200 rpm shaking incubators at 30°C, unless otherwise stated. Typically, cells were pre-cultured in YES or modified YNB over eight hours, followed by dilution to appropriate OD_595_ and sub-culture overnight to reach mid-exponential phase (OD_595_ 0.4-0.6) in the following morning. *S. japonicus* and *S. pombe* mating was induced on SPA solid medium containing supplements as above at 25°C. Spores were dissected and germinated on YES agar plates using a dissection microscope (MSM 400, Singer Instruments).

### Materials used in cell biological and biochemical experiments

D-glucose anhydrous (cat. #G/0450/60) and BD Difco™ Yeast Nitrogen Base without Amino Acids (cat. #291920) were purchased from ThermoFisher. Bacteriological agar (cat. #LP0011B) was purchased from Oxoid. Sodium phosphate dibasic dihydrate (cat. #71643), potassium hydrogen phthalate (cat. #P1088), adenine hemisulfate salt (cat. #A9126), L-histidine (cat. #H8000), L-leucine (cat. #L8000), uracil (cat. #U0750), sodium sulfate (cat. #239913), DMSO (cat. #D8418), terbinafine (cat. #T8826), amphotericin B (cat. #A2942), Tween 80 (cat. #P1754), filipin (cat. #F9765), 5-α-cholestane (cat. #C8003), ergosterol (cat. # E6510), squalene (cat. #S3626), tert-butyl methyl ether (cat. #650560), methanol (cat. #34860) and ethanol (cat. #32221) were purchased from Sigma-Aldrich. NuPage 4-12% BT gels (cat. #NP0321BOX), NuPage MOPS SDS running buffer (cat. #NP0001), NuPage transfer buffer (cat. #NP0006-1) and NuPage LDS sample buffer (cat. #NP0007) were purchased from Invitrogen. Nitrocellulose membranes 0.2 µm (cat. #1620112) were purchased from Bio-Rad. Mouse anti-RFP monoclonal antibody (cat. #6g6) was purchased from ChromoTek. Revert 700 Total Protein Stain Kit (cat. #926-11010) and IRDye 800CW Goat anti-Mouse IgG (cat. #926-32210) were purchased from LI-COR. RNeasy Plus Mini Kit (cat. #74134) and RNase-Free DNase Set (cat. #79256) were purchased from QIAGEN. Revertaid first strand cDNA synthesis kit (cat. #K1622) was purchased from Thermo Fisher. Probe Blue Mix Lo-ROX (cat. #PB20.21-01) was purchased from qPCRBIO. SPE Bulk Sorbent, primary secondary amine (PSA) (cat. #5982-8382) was purchased from Agilent. MSTFA (cat. #TS-48910) was purchased from Thermo Fisher and TSIM (cat. #MN701310.110) was purchased from Macherey-Nagel. The triterpenoid standards diplopterol (cat. #C1391.30), hop-17(21)-ene (cat. #C0789.30), hop-21(22)-ene (17β(H)) (cat. #C0699.30) and hop-22(29)-ene (cat. #C0698.30) were purchased from Chiron UK.

### Molecular genetics

All primers are shown in Data S3. Molecular genetics manipulations were performed using PCR^64^- or plasmid^65^-based homologous recombination. To express Shc1^S.j.^-sfGFP under the *tdh1* promoter in *S. pombe*, *shc1* open reading frame (ORF) was PCR-amplified from *S. japonicus* genomic DNA and cloned into pSO1006 (pAV0749^41^) between XhoI and EcoRI enzyme sites. The resulting plasmid pSO1275 was linearized before transformation and integrated into the *ura4* locus. To build plasmid pSO1276, *rga3* promoter (1203 bp upstream of the start codon) and *kanMX6* resistance cassette plus the plasmid backbone were PCR-amplified from *S. pombe* genomic DNA and pSO257 (pKS395) respectively, and assembled using the Gibson Assembly Master Mix (New England Biolabs). Plasmid pSO1276 was then used as a template to amplify *kanMX6:prga3* flanked by 80 base pairs upstream (position -529) and downstream (position +1) of the endogenous *erg1* promoter, followed by integration into the endogenous locus, resulting in the replacement of *S. pombe erg1* endogenous promoter by a weaker *rga3* promoter. A PCR-based method was used to knock out *S. japonicus shc1* and *erg1*, as well as to tag Shc1, and *S. pombe* and *S. japonicus* Erg1 at the C-terminus using *kanR* or *natR* as selection markers. All constructs were verified by sequencing. *S. japonicus* cells were transformed by electroporation^62^. *S. pombe* transformation was performed using lithium acetate and heat shock^60^. Transformants were selected on YES agar plates containing G418 (Sigma Aldrich), nourseothricin (HKI Jena), or EMM agar plates minus uracil.

### Serial dilutions assays

*S. japonicus* and *S. pombe* cells were pre-cultured overnight in YES or modified YNB at 30°C until early-exponential phase. Cultures were then diluted to OD_595_ 0.1, and serial 10-fold dilutions were spotted on YES or YNB agar plates or the same media supplemented with different concentrations of terbinafine (Sigma-Aldrich) or amphotericin B (Sigma-Aldrich). YES plates supplemented with Tween 80 (Sigma-Aldrich) were made by adding 1% of a Tween 80 stock solution prepared with pure ethanol as solvent (50% v/v). Plates were typically incubated at 30°C unless stated otherwise, either in the presence of oxygen or in an anoxic environment inside an InvivO_2_ 400 workstation (Baker-Ruskinn). After three days, plates were scanned using an Epson Perfection V700 Photo scanner. All experiments were repeated three times with similar results, and representative experiments are shown in the corresponding figures.

### Microscopy and image analysis

Prior to imaging, 1ml cell culture was concentrated to 50 μl by centrifugation at 1500 x g for 1 min. 2 μl cell suspension was loaded under a 22 × 22 mm glass coverslip (VWR, thickness: 1.5). Fluorescence images in Figures 1b, 4a and 4g were acquired using Yokogawa CSU-X1 spinning disk confocal system with Eclipse Ti-E Inverted microscope with Nikon CFI Plan Apo Lambda 100× Oil N.A.=1.45 oil objective, 600 series SS 488nm SS 561nm lasers and Andor iXon Ultra U3-888-BV monochrome EMCCD camera controlled by Andor IQ3. Single plane images with inverted LUT (look-up-table) are shown. Images shown in Figure 1e and S1a were captured using a Zeiss Axio Observer Z1 fluorescence microscope fitted with α Plan-FLUAR 100×/1.45 NA oil objective lens (Carl Zeiss) and the Orca-Flash4.0 C11440 camera (Hamamatsu). Images were taken at the medial focal plane of cells. Filipin staining of sterols was performed by adding the drug at a final concentration of 5 µg/ml from a DMSO stock to the cell liquid cultures in YES medium. Cells were observed immediately upon drug addition. Image analysis and quantification were performed using Fiji^66^. Within the same experiment, images are directly comparable as they are adjusted to equal brightness and contrast levels. Measurements of cellular length and width were performed on bright-field images acquired with the Zeiss epifluorescence microscope.

### Cell growth and colony forming unit (CFU) assays

For growth rate experiments, *S. japonicus* and *S. pombe* cells were grown overnight in YES or modified YNB either at 30°C, 36°C or 37°C until early-exponential phase. Cultures were then diluted to OD_595_ 0.1-0.15 with the same medium and loaded into a 96-well plate. Growth was measured every 60 min at the corresponding temperature using VICTOR Nivo multimode plate reader (PerkinElmer). Growth rates and Tmid were calculated using the Growthcurver R package^67^. Experiments were repeated at least three times from cultures grown on separate occasions.

CFU measurements were performed as described in^68^ with minor modifications. Briefly, *S. japonicus* cells were pre-cultured overnight in YES or modified YNB at 30°C until stationary phase. Cultures were normalized to OD_595_ 1 and three decimal dilutions (10^-1^, 10^-2^, 10^-3^) of each were prepared. 100 µl of the 10^-3^ dilution were further diluted with 500 µl of the appropriate media, and 100 µl of the resultant sample were plated in quadruplicates on YES or modified YNB agar plates. After two days (YES plates) or three days (modified YNB plates) plates were scanned using an Epson Perfection V700 Photo scanner. The colony numbers and size were measured using Fiji^66^.

### Western blotting of Erg1-mCherry

*S. japonicus* and *S. pombe* cell cultures were grown in YES to mid-exponential phase and 5 ODs were pelleted for 1 min at 2103 x g and liquid medium was removed. Cells were resuspended in 1 ml ice-cold dH_2_O and transferred to 1.5 ml Eppendorf tubes. Cells were washed and snap frozen in liquid nitrogen. Western blotting experiments were performed as previously described in^69^ with minor modifications. For Erg1-mCherry detection nitrocellulose 0.2 µm membranes (Bio-Rad) were blocked for 1 h with TBST buffer containing 5% skimmed milk and then incubated for 1 h with mouse α-RFP antibody (ChromoTek). Membranes were washed with TBST and incubated for 1 h in TBST buffer with IRDye800 conjugated α-mouse antibody (LI-COR Biosciences). Total protein levels were detected using the LI-COR Revert 700 Total Protein stain kit (LI-COR Biosciences). Proteins were detected using the Odyssey Infrared Imaging System (LI-COR Biosciences). Samples were collected as at least three biological replicates from cultures grown on separate occasions. Quantification was performed using Fiji^66^.

### Reverse transcription and real-time quantitative PCR (RT-qPCR)

Reverse transcription and qPCR were performed as previously described in^29^. The RT-qPCR was performed on a LightCycler 96 Instrument (Roche Diagnostics) in three biological and two technical repeats. RT-qPCR signal was normalized to actin expression levels.

### Triterpenoid detection by GC-MS

*S. japonicus* and *S. pombe* cell cultures were grown in modified YNB to mid-exponential phase and 10 ODs were harvested via filtration, then snap frozen in liquid nitrogen. For triterpenoid detection, a total of five replicates were collected per condition (three biological repeats, two technical repeats). Metabolite extraction for triterpenoid detection was performed as described in ^6,70^ with minor modifications. Before processing, cell pellets were lyophilized overnight using a freeze dryer lyophilizer (Labconco). For cell lysis, each freeze-dried pellet was mixed with 1 ml of 2M NaOH, transferred to glass vials and heated for 1h at 70°C in a water bath, with vortexing at 15 min intervals. After saponification, suspensions were allowed to cool to room temperature and divided into two microcentrifuge tubes (2 x 500 µl). 650 μl of distilled methyl-tert-butylether (MtBE, Sigma-Aldrich, HPLC grade) and 100 μl of internal standard solution (5α-cholestane in MtBE, 10 μg/ml) were added to each sample. Mixtures were vortexed for 1 min and centrifuged at 9000 x g for 5 min. After centrifugation, the organic upper layer (∼550 µl) was transferred into a new microcentrifuge tube containing 40 ± 2 mg of a mixture (7:1) of anhydrous sodium sulfate (Sigma-Aldrich) and primary secondary amine (PSA, Agilent Technologies) (dispersive solid phase). Lysed cells were subjected to a second round of metabolite extraction by adding another 750 μl of MtBE, shaking for 1 min and centrifuging at 9000 x g for 5 min. The organic upper layer (∼650 µl) was transferred to the tube containing the dispersive solid phase and combined with the previous organic extract. The mixtures were shaken and centrifuged again, and 1 ml of the purified upper layer was transferred to an amber glass vial (Agilent Technologies).

Samples were evaporated to dryness under a stream of nitrogen at room temperature. For each pair of vials, one residue was resuspended in 700 μl of MtBE and 50 μl of silylation reagent mixture MSTFA/TSIM (9:1) and the other residue was resuspended in 750 μl of MtBE. Samples were vortexed for 10 sec and incubated for at least 30 min at room temperature.

Triterpenoid analysis was performed by GC-MS using an Agilent 7890B-5977A system. Splitless 1 µl injection (injection temperature 250°C) onto a 40 m × 0.25 mm VF-5ms + EZ Guard column (Agilent J&W) was used with helium as the carrier gas, in electron impact (EI) ionization mode. The initial oven temperature was 55°C (1 min), followed by temperature gradients to 260°C at 50°C/min, and then to 320°C at 4°C/min (held for 4 min). Mass spectra were acquired at 70 eV in the range from 50 m/z to 600 m/z. Triterpenoid identification was performed by comparison to retention time and fragment ion pattern of authentic standards using MassHunter Workstation software (B.06.00 SP01, Agilent Technologies) and confirmed by comparison to deconvoluted mass spectra of those in the NIST Mass Spectral Library software (NIST 23, software version 3.0). The following standards we used: diplopterol, ergosterol, hop-17(21)-ene, hop-21(22)-ene (17β(H)), hop-22(29)-ene and squalene. A 6-point calibration curve of the GC-MS system with standard mixtures was used for relative quantification of triterpenoid compounds.

### ESI-MS-based lipidomic analysis

Lipid standards were from Avanti Polar Lipids (Alabaster, AL, USA). Solvents for extraction and MS analyses were liquid chromatography grade (Merck, Darmstadt, Germany) and Optima LC-MS grade (Thermo Fisher Scientific, Waltham, MA, USA), as applicable. All other chemicals were the best available grade purchased from Sigma-Aldrich or Thermo Fisher Scientific.

Exponentially growing yeast cell cultures in modified YNB were pelleted. For lipidomics analysis, a total of five replicates were collected per condition (three biological repeats, two technical repeats). Cellular lipids were extracted using previously described methods^29^. Lipidomics data are expressed as mol% of polar lipids; polar lipids include all measured lipids except DG, TG, EE, and sterols. Double bond index (DBI), average chain length and average lipid species profile was calculated for the sum of major GPLs (PC, PI, PE and PS). DBI was calculated as Σ(db x [GPLi])/Σ[GPLi], where db is the total number of double bonds in fatty acyls in a given GPL species, and the square bracket indicates mol% of GPLs. Average chain length was calculated as Σ(C x [GPLi])/Σ[GPLi], where C is the total number of carbons in fatty acyls in a given GPL, and the square bracket indicates mol% of GPLs.

### Materials used in biophysical experiments

DMSO (cat. #D8418), D-glucose (cat. #G/0450/60), DPH (cat. #D208000), EDTA (cat. #E5134), ergosterol (cat. #E6510), HEPES (cat. #H3375), NaCl (cat. #S7653), Na_2_HPO_4_.2H_2_O (cat. #71643), NaH_2_PO_4_.H_2_O (cat. #71507) were purchased from Sigma-Aldrich. Sucrose (cat. #F492423) was purchased from Fluorochem. FAST DiI (cat. #D7756) was purchased from Thermo Fisher Scientific and C-laurdan (cat.#7273) was purchased from Tocris Bioscience. Synthetic glycerophospholipids including POPC (850457C), DPPC (850355C) and DOPC (850375C) were purchased from Avanti Polar Lipids. 18:0/10:0 PC (SDPC) was custom-synthesized by Avanti Polar Lipids. 17Beta(H),21Beta(H)-22-Hydroxyhopane (cat #C1391.30) commonly known as diplopterol was purchased from Chiron UK. Spectroscopic grade solvents such as methanol (cat. #154903) and chloroform (cat. #366919) used for lipid stock preparation were purchased from Sigma-Aldrich. Milli-Q water was used throughout. The concentration of a stock solution of DPH in methanol was estimated from its molar extinction coefficient (€) of 88,000 M^-1^cm^-1^ at 350 nm^71^. C-laurdan stock solution (100 µM) was prepared in DMSO.

### Preparation of Giant Unilamellar Vesicles (GUVs)

GUVs were prepared by electroformation using the Nanion Vesicle Prep Pro (Nanion Technologies, Munich, Germany) as described previously^72,73^, with minor modifications. Lipids (200 nmol for two-component liposomes or 300 nmol for three-component liposomes) along with 0.2 mol% FAST DiI probe were dried onto the conductive side of the ITO-coated slide. The glass slide was dried in vacuum for ∼2.5h. A medium O-ring was coated with vacuum grease and placed around the dried lipid film. 270 µl of 250 mM sucrose was added inside the O-ring, and the second conductive slide was placed on the top, making a sandwich. The standard protocol for vesicle preparation of 120 min was used with a rise and fall of 3 min. The amplitude was set at 10 Hz, voltage was 3 V, and the temperature was 55°C. For C-laurdan labelling, the probe was added to electroformed GUVs from a 100 µM stock solution in DMSO such that the final lipid-to-probe ratio was 300:1 (mol/mol). GUVs were stored at 4°C and used for imaging measurements on the following day. For spinning disk confocal microscopy measurements, 20 µl of GUV solution was slowly added to a microscope chamber (ibidi, Gräfelfing, Germany) filled with ∼400 µl of 250 mM glucose solution for 2h. This allowed vesicles to settle onto the bottom of the chamber before imaging measurements were carried out.

### Preparation of Large Unilamellar Vesicles (LUVs)

LUVs were prepared as described previously with slight modifications^74^. For fluorescence anisotropy measurements, 300 nmol lipids and 3 nmol DPH (lipid-to-probe ratio 300:1) were mixed and dried using a nitrogen stream while being warmed gently at 37°C. For cryo-EM measurements, liposomes were prepared using 600 nmol of lipids. The lipid samples were dried in vacuum for 3 h, followed by hydration at 60°C for 1h in 1 ml of buffer A (10 mM sodium phosphate, 150 mM NaCl, pH 7.4) for fluorescence anisotropy measurements or buffer B (10 mM HEPES, 150 mM NaCl, pH 7.4) for cryo-EM. Samples were vortexed for 1 min to form homogeneous multilamellar vesicles. LUVs were prepared by extrusion using Avestin Liposofast Extruder (Ottawa, Canada) as previously described^75^. Briefly, multilamellar vesicles were freeze-thawed five times using liquid nitrogen and extruded through polycarbonate filters (pore diameter of 100 nm) mounted on extruder fitted with Hamilton syringes (Hamilton Company, Reno, NV). The samples were subjected to uneven passes (21 passes for all liposomes except the gel-like liposomes with DPPC and ergosterol, for which 33 passes were used) on a 55°C hot plate. LUV sizes were measured using dynamic light scattering (Malvern Zetasizer Nano ZS). For anisotropy experiments, samples were kept overnight at 25°C before measurements. For cryo-EM experiments LUVs were stored at 4°C overnight before preparing samples.

### Water permeability measurements

For water permeability assays, unlabelled single or two-component MLVs were prepared in Tris–HCl 5 mM, pH 7.0, 100 mM sucrose (isotonic buffer)^76^. Isotonic MLV suspensions were diluted by adding 0.12 ml of MLVs to 2 ml of hypotonic buffer (Tris– HCl 5 mM, pH 7.0) equivalent to ∼16 times dilution. The time-dependent reduction in absorbance at 550 nm (Fig. S2c) due to loss in turbidity was monitored for 6 min, and used to calculate permeability coefficients (Fig. S2d). Absorbance was measured using V-560 spectrophotometer (Jasco, West Yorkshire, UK) at room temperature (∼25°C).

### Visualizing phase separation

Images of FAST DiI-labelled GUVs were obtained using Yokogawa CSU-X1 spinning disk confocal system mounted on the Eclipse Ti-E Inverted microscope with Nikon CFI Plan Apo Lambda 100X Oil N.A. = 1.45 oil objective, 600 series SS 488nm, SS 561nm lasers and Andor iXon Ultra U3-888-BV monochrome EMCCD camera. Excitation laser of 561 nm was used. Z-stacks were acquired with step size of 0.6µm for 12 steps. Imaging was performed at 25°C. Image processing and quantifications were performed in Fiji^66^. Fluorescence images are shown with inverted LUT (look-up table) (Fig. 2b, c, and Fig S2 b, e).

### Confocal spectral imaging for estimating membrane order

Spectral imaging of GUVs was performed on a Zeiss LSM 880 confocal microscope equipped with a 32-channel GaAsP detector array, as described previously^77^. Excitation wavelength of 405 nm was used for fluorescence excitation of C-laurdan while the lambda detection range was set between 415 and 691 nm. The intervals between the individual detection channels were set to 8.9nm which allowed the simultaneous coverage of the whole spectrum with 32 detection channels. The .czi format images were analyzed using a custom GP plug-in compatible with Fiji/ImageJ^77^. The fluorescence intensities obtained after processing the confocal images were used to calculate GP (generalized polarization) according to the equation:

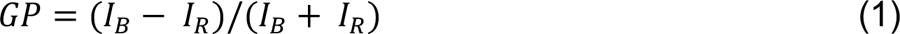

where *I*_B_ and *I*_R_ are the fluorescence intensities corresponding to the blue-shifted wavelength (442 nm) and the red-shifted wavelength (496 nm) for C-laurdan (Fig. 2e). Confocal imaging measurements were carried out at 37°C.

### Fluorescence anisotropy measurements in liposomes

Steady-state fluorescence anisotropy measurements were performed at 25°C on a FP-8500 spectrofluorometer (Jasco, West Yorkshire, UK) with in-built excitation and emission polarizers. For monitoring DPH fluorescence, the excitation wavelength was set at 358 nm and emission was monitored at 430 nm. Quartz cuvettes with a path length of 1 cm were used. Excitation and emission slits with bandpass of 5 and 10 nm were used for all measurements. Fluorescence was monitored with a 30 s interval between successive openings of the excitation shutter to reverse any photoisomerization of DPH^78^. Anisotropy values were obtained simultaneously along with the measurements according to the equation^79^:

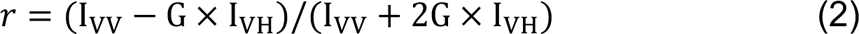

where I_VV_ and I_VH_ are the fluorescence intensities measured with the excitation polarizer oriented vertically and the emission polarizer oriented vertically and horizontally oriented, respectively. G is the grating factor and is the ratio of the efficiencies of the detection system for vertically and horizontally polarized light, and is equal to I_HV_/I_HH_. G factor was measured before sample measurements and was always ∼1.

### Cryo-EM grid preparation, imaging and data processing

A thin carbon film was deposited onto a mica sheet using EMITECH K950X, and transferred onto Quantifoil (200 Cu mesh, R2/2) grids. The carbon film grids were then glow discharged (EMITECH K100X, 25 mA, 30 s) in an amylamine atmosphere. 4 μl of sample was pipetted to the grid in the environmental chamber of a Vitrobot Mark IV (FEI/Thermo) at 4°C and 95-100% humidity. The grid was blotted for 1.5 s before plunging into liquid ethane kept at liquid nitrogen temperature. The grids were imaged on a Talos Arctica microscope (FEI/Thermo) at 200 kV using EPU software (v 2.11). Movies were recorded on a Falcon III camera in linear mode with a total dose of 48 electrons per Å^2^ fractionated over 10 frames (dose rate 24 e^-^/Å^2^/s) with a 1.99 Å pixel size and a nominal defocus of -2 μm. All movies were imported into Relion (v 4.0.0)^80^, followed by Relion’s own motion correction and CTF estimation (CTFFIND, v 4.1.13)^81^. The images within the defocus range from -1.5 μm to -2.1 μm were selected, then converted to 16-bit tiff format for further measurements. Subsequent analysis of images was performed using Fiji/Image. A 2-point Gaussian filter was applied for clear distinction of dips in the intensity profile before computation of D_TT_ ^38^ from the images.

### Statistics and reproducibility

The statistical details of experiments, including the number of biological and technical replicates and the dispersion and precision measures can be found in Figure Legends and Materials and Methods. All data were analysed using unpaired t-test statistical analysis, unless indicated otherwise. All plots were generated using GraphPad Prism 10.

No data were excluded in the cell biological and physiological experiments. In the measurements of triterpenoid abundance, linear regression analyses between sample OD_595_ and lipid content were performed for each experimental group. Samples showing anomalous lipid levels relative to the sample amount were excluded as likely sample preparation artifacts.

## Acknowledgements

We are grateful to the Oliferenko lab for discussions and Eugene Makeyev for suggestions on the manuscript. Many thanks to Fred Betts Thompson for media preparations and Nikita Komarov for helping with *shc1Δ* mutant construction, and to the Crick Metabolomics and Structural Biology STPs for invaluable training and assistance. Elisa Gomez Gil is supported through a long-term EMBO postdoctoral fellowship (ALTF 712-2022). We thank the Single Cell Omics Advanced Core Facility staff of the HCEMM and Biological Research Center for help with their resources and their support. HCEMM has received funding from the EU’s Horizon 2020 research and innovation program under grant agreement No. 739593 and KIM NKFIA 2022-2.1.1-NL-2022-00005. This work was supported by the Francis Crick Institute, which receives its core funding from Cancer Research UK (CC0102), the UK Medical Research Council (CC0102), and the Wellcome Trust (CC0102), and the Wellcome Trust Senior Investigator Award (103741/Z/14/Z) and Wellcome Trust Investigator Award in Science (220790/Z/20/Z) to Snezhana Oliferenko.

## Author contributions

B. D. R. conceived and performed biophysical experiments; analyzed data; and co-wrote the manuscript. E. G. G. conceived and performed cell biological and biochemical experiments; generated strains; analyzed data; and co-wrote the manuscript. M. P. and G. B. designed, performed, and interpreted all ESI-MS lipidomics experiments and edited the manuscript. V. N. and J. I. M. designed, performed and interpreted GC-MS experiments and edited the manuscript. Q. C. and P. R. designed, performed and interpreted cryo-EM experiments and edited the manuscript. S. O. conceived and interpreted experiments and co-wrote the manuscript. This research was funded in whole, or in part, by the Wellcome Trust (103741/Z/14/Z; 220790/Z/20/Z to S. O.). For the purpose of Open Access, the author has applied a CC-BY public copyright licence to any Author Accepted Manuscript version arising from this submission.

## Competing interests

The authors declare no competing or financial interests.

**Data S1. Lipidomics. All data presented as Excel tables.** (i) Detailed lipidomics dataset for comparison of *S. japonicus wild type* (*WT*), *shc1Δ* and *erg1Δ* cells and *WT* cells in the absence of oxygen. Whole lipidome species composition is shown. (ii) Detailed lipidomics dataset for comparison of *S. pombe WT* and *ptdh1:shc1^S.j.^-sfGFP* cells. Whole lipidome species composition is shown. Data are expressed as mol% of polar lipids; polar lipids include all measured lipids except DG, TG, EE, and sterols. Sterols were analyzed by GC-MS, all other lipids by shotgun ESI-MS. Data are expressed as average ± S.D., n= 5 (three biological and two technical replicates). Student’s t-tests were performed for pairwise multiple comparisons; significance was accepted for p < 0.05. Blue fill indicates p < 0.05. PC, phosphatidylcholine; PE, phosphatidylethanolamine; MMPE, monomethyl PE; DMPE, dimethyl-PE; PI, phosphatidylinositol; PS, phosphatidylserine; PG, phosphatidylglycerol; PA, phosphatidic acid; CL, cardiolipin; LPC, LPE, LPI, LPS, LPA, LCL, the corresponding lyso lipids; Cer, ceramide; IPC, inositolphosphoceramide; MIPC, mannosyl-inositolphosphoceramide; DG, diacylglycerol; TG, triacylglycerol; EE, ergosteryl ester. DBI, double bond index; Av_ChL, average chain length. Sum formulas for glycero(phospho)lipids (GPL) are defined as the lipid class abbreviation followed by the total number of carbons and total number of double bonds for all chains, e.g. PC(34:1). Sum formulas for sphingolipids are defined as the lipid class abbreviation followed by the total number of carbons, total number of double bonds and total number of hydroxyl groups in the long chain base and the fatty acyl moiety, e.g. Cer(44:0:3). Double bond index (DBI) was calculated for GPLs (PC, PE, PI and PS) as Σ(db x [GPLi])/Σ[GPLi], where db is the total number of double bonds in fatty acyls in a given GPL species, and the square bracket indicates mol% of GPLs. Average acyl chain length (Av_ChL) was calculated for GPLs (PC, PE, PI and PS) as Σ(C x [GPLi])/Σ[GPLi], where C is the total number of carbons in fatty acyls in a given GPL, and the square bracket indicates mol% of GPLs. Average species profile was calculated for PC, PE, PI and PS. SoamD score was calculated for all lipid species as Σabs(mol%_mutant_ – mol%*_WT_*). Please note that the list of lipid species is matched for the two yeast species. Due to the substantial difference in their lipid profile, one can find zero values for certain lipid species in *S. japonicus* or *S. pombe*. (iii) Detailed triterpenoid abundance dataset for comparison of *S. pombe WT*, *ptdh1:shc1^S.j.^-sfGFP*, *prga3:erg1* and *ptdh1:shc1^S.j.^-sfGFP prga3:erg1* cells analyzed by GC-MS. Data are expressed µM/OD, average ± S.D., n= 4 (two biological and two technical replicates). (iv) Shotgun lipidomics quantification details. (v) GC-MS quantification details.

**Data S2. List of fission yeast strains used in this study.** *S. japonicus* and *S. pombe* strains used in each Figure are listed in individual spreadsheets in order of appearance.

**Data S3. List of primers used in this study.** Primers used to generate deletions, fluorescent protein tagging and promoter replacement, and genotyping primers are listed in individual spreadsheets.

---

**Figure S1.**
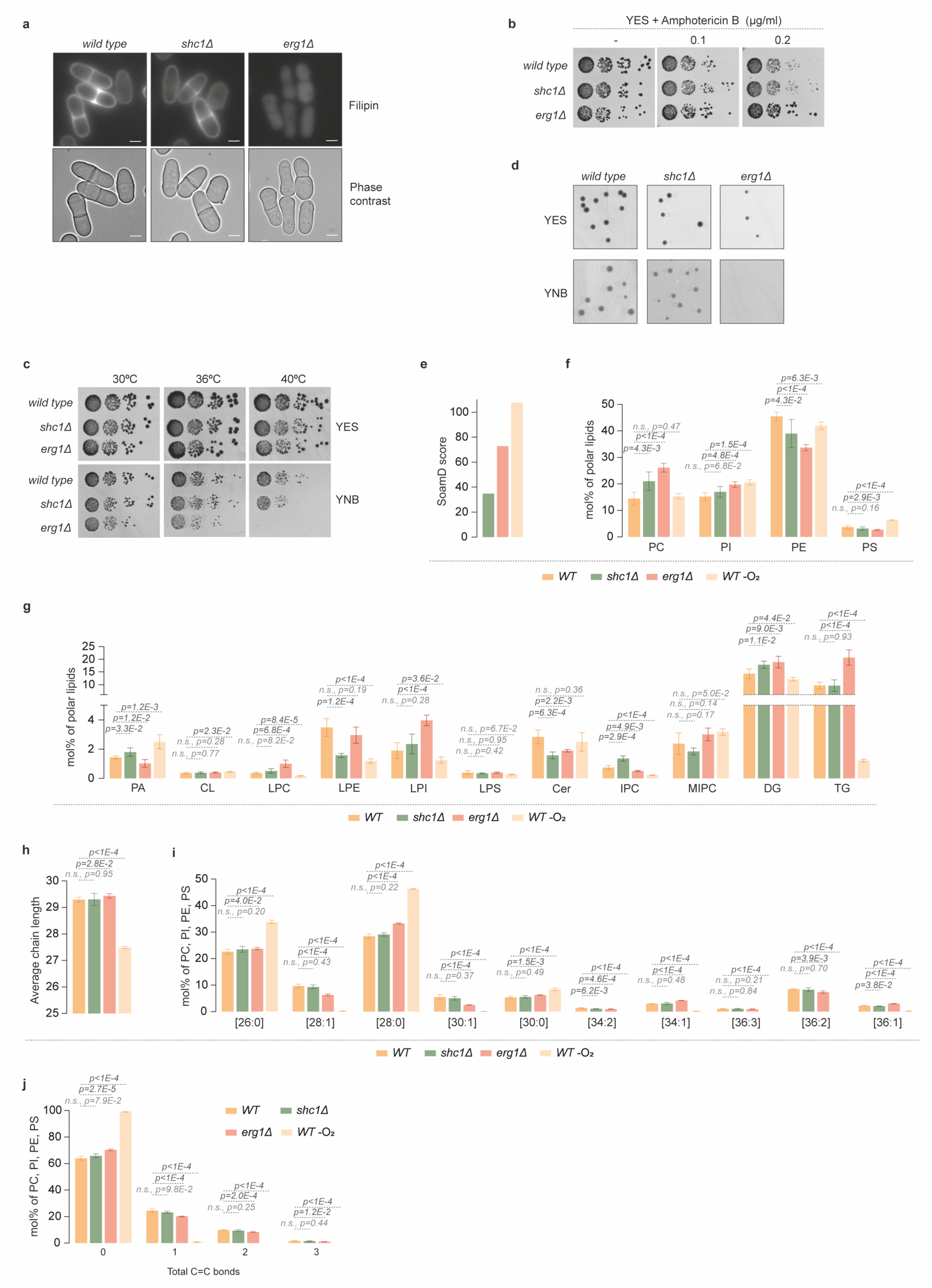
Related to Figure 1. *S. japonicus* lipidome responds to the lack of either ergosterol or hopanoids. (**a**) Micrographs of filipin-stained *S. japonicus wild type* (*WT*), *shc1Δ* and *erg1Δ* cells grown in YES medium. Brightfield images are also included. Scale bars represent 5 µm. (**b**) Serial dilution assay of *S. japonicus* strains of indicated genotypes carried out in YES medium in the absence or the presence of amphotericin B. (**c**) Serial dilution assay of *S. japonicus* strains of indicated genotypes performed in indicated conditions. (**d**) Representative images of colonies from the CFU assay shown in Fig. 1i, j. (**e**) Sum of absolute mol% difference (SoamD) relative to *S. japonicus WT* for every lipid species. (**f**) Relative abundance of the four main GPL classes (PC, PI, PE and PS) in *S. japonicus WT*, *shc1Δ* and *erg1Δ* cells grown in normoxia, and *WT* cells grown in anoxia. (**g**) Relative abundance of the indicated lipid classes: phosphatidic acid (PA), cardiolipin (CL), lysophosphatidylcholine (LPC), lysophosphatidylethanolamine (LPE), lysophosphatidylinositol (LPI), lysophosphatidylserine (LPS), ceramide (Cer), inositol phosphoceramide (IPC), mannosyl-inositolphosphoceramide (MIPC), diacylglycerol (DG) and triacylglycerol (TG) in *S. japonicus WT*, *shc1Δ* and *erg1Δ* cells grown in normoxia, and *WT* cells grown in anoxia. (**h**) Average combined FA length calculated for the sum of PC, PI, PE, and PS in *S. japonicus* strains of indicated genotypes and conditions. (**i**) Molecular species composition calculated for the sum of PC, PI, PE, and PS in *S. japonicus WT*, *shc1Δ* and *erg1Δ* cells grown in normoxia, and *WT* cells grown in anoxia. The categories are shown as the total number of carbon atoms: total number of double bonds in acyl chains. (**j**) Grouping of GPL species according to the number of double bonds calculated for the sum of PC, PI, PE, and PS in *S. japonicus WT*, *shc1Δ* and *erg1Δ* cells grown in normoxia, and *WT* cells grown in anoxia. (**f**-**j**) Data are represented as average ± S.D (n=5). p-values are derived from two-tailed unpaired t-test.

**Figure S2.**
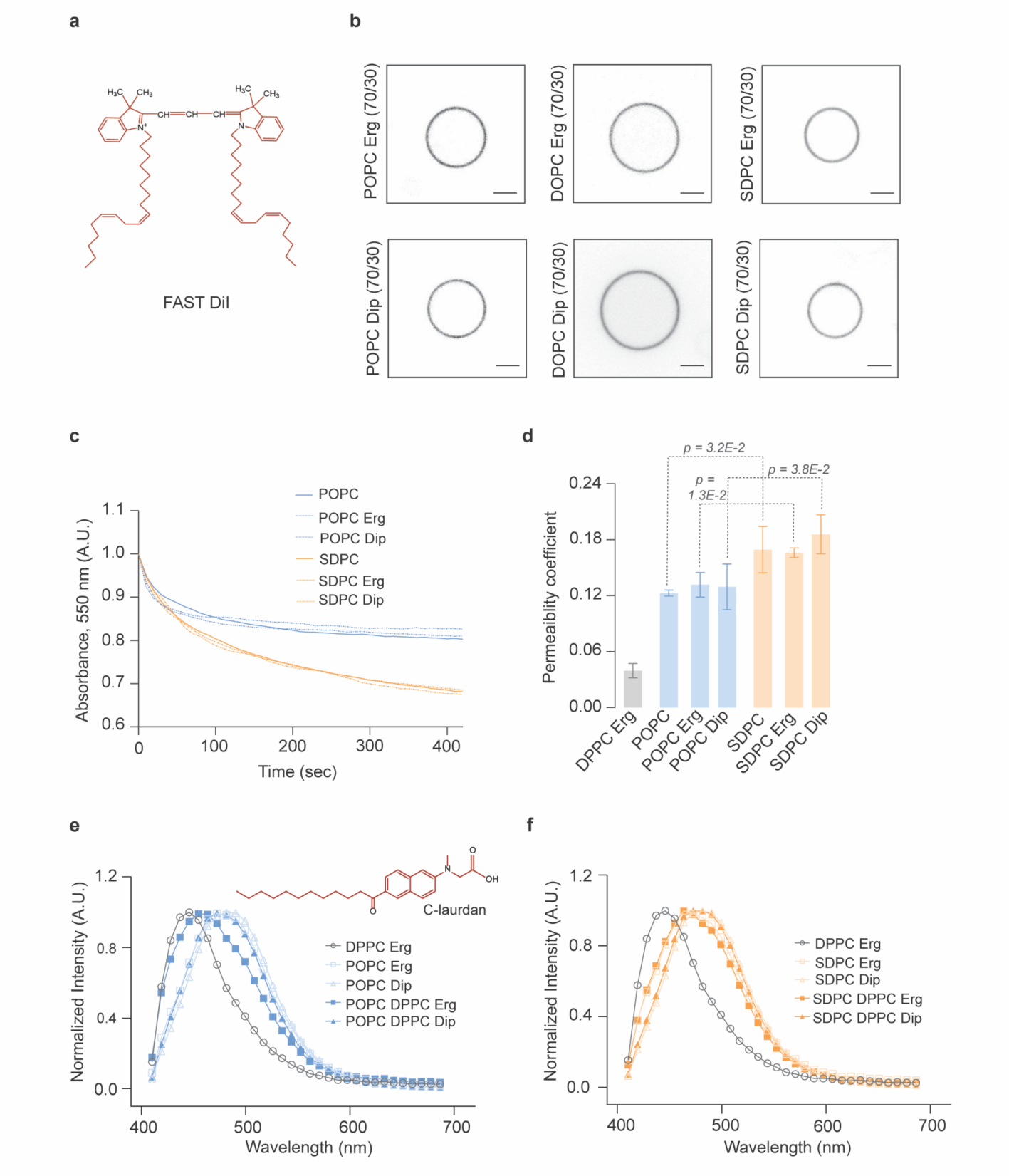
Related to Figure 2. Exploration of biophysical properties of model membranes containing asymmetrical saturated or symmetrical unsaturated glycerophospholipids. (**a**) Chemical structure of the probe 1,1’-Dilinoleyl-3,3,3’,3’-Tetramethylindocarbocyanine, 4-Chlorobenzenesulfonate (FAST DiI) used for labelling GUVs in phase-separation experiments. (**b**) Representative spinning disk confocal images of two-component liposomes. Incorporation of FAST DiI into disordered membranes assembled from either unsaturated symmetrical lipids (POPC or DOPC) or the saturated asymmetrical lipid (SDPC) with either 30 mol% ergosterol (*top panel*) or diplopterol (*bottom panel*). Mid-planes from confocal z-stacks are shown, scale bars represent 2 μm. (**c**) Time-dependent decrease in absorbance measured in liposomes. The representative curves for time-dependent reduction in absorbance is shown in single and two-component MLVs made with POPC or SDPC with either 30 mol% ergosterol or diplopterol. (**d**) Permeability coefficients estimated from (**c**) (average ± S.D. from three independent measurements). (**e**, **f**) Representative normalized intensity vs wavelength curves obtained from C-laurdan (structure shown in inset) imaging in two- and three-component GUVs made with the unsaturated symmetrical glycerophospholipid POPC (**e**) or saturated asymmetrical SDPC (**f**). Data for gel-like membranes made with DPPC and ergosterol is shown for comparison.

**Figure S3.**
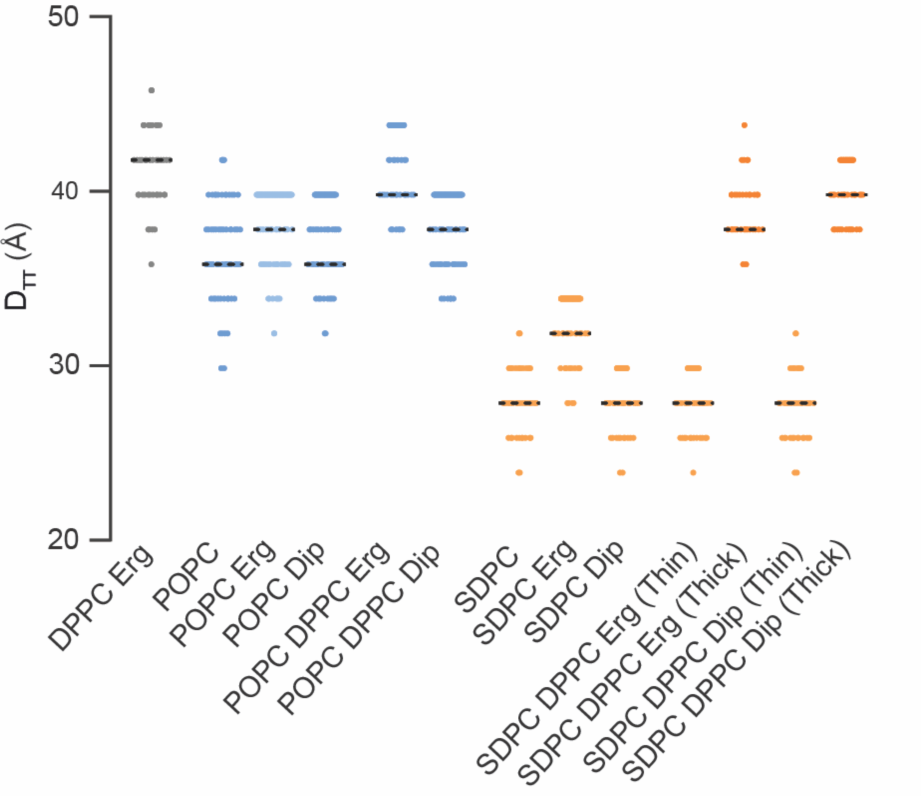
Related to Figure 3. Estimation of D_TT_ from cryo-EM measurements in liposomes. Distribution of D_TT_ values from individual measurements in single, two- and three-component LUVs made with either symmetrical unsaturated POPC or asymmetrical saturated SDPC. Specific LUV compositions are indicated.

**Figure S4.**
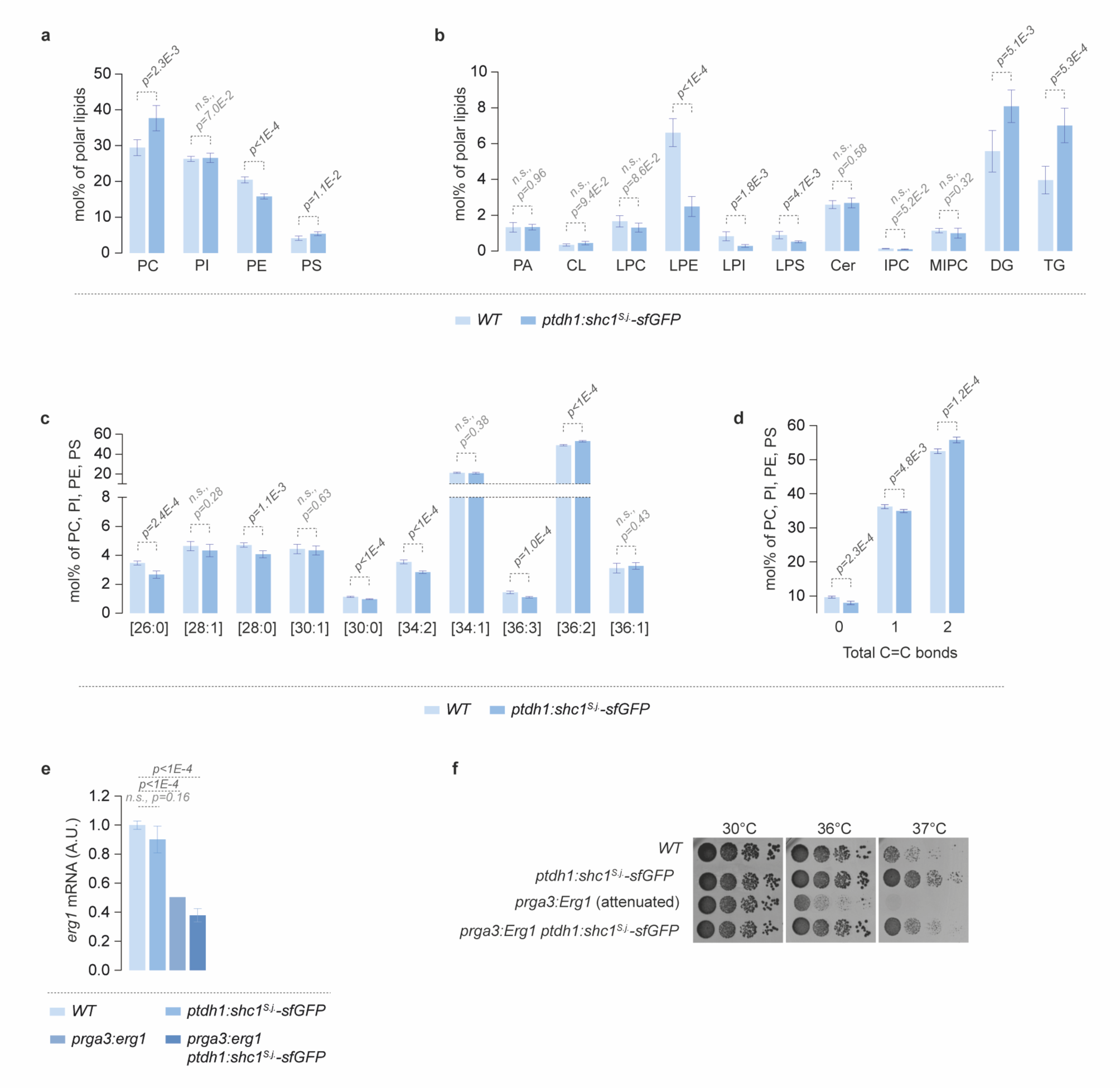
Related to Figure 4. *S. pombe* lipidome adapts to the presence of hopanoids. (**a**) Relative abundance of the four main GPL classes (PC, PI, PE and PS) in *S. pombe* cells of indicated genotypes. (**b**) Relative abundance of the indicated lipid classes (PA, CL, LPC, LPE, LPI, LPS, Cer, IPC, MIPC, DG and TG) in *S. pombe wild type* (*WT*) and *ptdh1:shc1^S.j.^-sfGFP* cells. (**c**) Molecular species composition calculated for the sum of PC, PI, PE, and PS in *S. pombe WT* and *ptdh1:shc1^S.j.^-sfGFP* cells. The categories are shown as the total number of carbon atoms: total number of double bonds in acyl chains. (**d**) Grouping of GPL species according to the number of double bonds calculated for the sum of PC, PI, PE, and PS in *S. pombe WT* and *ptdh1:shc1^S.j.^-sfGFP* cells. (**e**) Steady-state *erg1* mRNA levels in *S. pombe* strains of indicated genotypes grown in YES, as measured by qPCR, normalized to the *WT*. Results are shown as average ± S.D. (n=3). (**f**) Serial dilution assay of *S. pombe* strains of indicated genotypes carried out at indicated temperatures in YES medium. (**a**-**d**) Data are represented as average ± S.D. (n=5). (**a**-**e**) p-values are derived from two-tailed unpaired t-test.

